# Competition for microtubule lattice spacing between a microtubule expander and compactor

**DOI:** 10.1101/2025.02.28.640185

**Authors:** Alexandra L. Paquette, Sofía Cruz Tetlalmatzi, Justin A. G. Haineault, Adam G. Hendricks, Gary J. Brouhard, Muriel Sébastien

**Author notes:** These authors participated equally to this work (Alexandra L. Paquette, Sofía Cruz Tetlalmatzi). Correspondence should be addressed to M.S. and G.J.B.

## Abstract

Microtubules exist in expanded and compacted states, as defined by the lattice spacing of αβ-tubulin dimers. Changes in lattice spacing has been linked to factors such as GTP-hydrolysis, the binding of microtubule-associated proteins (MAPs), the tubulin code, and microtubule bending. These diverse factors exert opposing molecular driving forces on the microtubule lattice that push lattice spacing towards expanded or compacted states. To better understand how these opposing forces are reconciled, we developed in vitro and cell-based model systems for the competition between a microtubule expander (paclitaxel) and a microtubule compactor (Doublecortin, or DCX). Using an in vitro reconstitution approach, we show that paclitaxel expands microtubules cooperatively. In cells, high concentrations of paclitaxel cause DCX to relocalize to compacted lattices found at concave bends. When the concentration of DCX is increased, however, we find that DCX re-compacts the previously expanded microtubules in vitro. Consistently, high expression levels of DCX prevent its relocalization in paclitaxel treated cells. When the competition between paclitaxel and DCX is “balanced”, we observe a complex phenotype: DCX simultaneously localized to both long, straight clusters and concave bends, while other regions on the microtubule network remained DCX-free. We conclude that multiple lattice spacings can coexist in cells. Our results indicate that competition for microtubule lattice spacing is a critical aspect of microtubule physiology.

## Introduction

Microtubules are cylindrical lattices of αβ-tubulin dimers, and these lattices exist in diverse states due to their intrinsic structural plasticity (Kueh and Mitchison 2009). A key structural feature of the microtubule lattice is its “lattice spacing”, also known as its “dimer rise”, defined here as the length of a single αβ-tubulin dimer along the long axis of its protofilament. Tubulin is a GTPase, and differences in lattice spacing were observed between different nucleotide states in early cryo-electron microscopy (cryo-EM) (Hyman *et al*. 1995; Alushin *et al*. 2014; Zhang *et al*. 2015). Measurements of lattice spacing cluster into two states: (1) an “expanded” state with a lattice spacing of 83.5 ± 0.2 Å and (2) a “compacted” state with a lattice spacing of 81.7 ± 0.1 Å, a difference of 2.3 ± 0.2 % (mean ± SEM, (LaFrance *et al*. 2022)). The “accordion-like” transitions between these two states reorganize the contact surfaces of tubulin-tubulin bonds (Manka and Moores 2018b). In cells, expanded and compacted lattices have been observed adjacent to one another by cryo-electron tomography (de Jager *et al*. 2024), and changes in lattice spacing occur in response to osmotic shifts (Shen and Ori-McKenney 2024). These observations suggest that lattice spacing may be regulated *in vivo*. However, the implications of these two lattice spacings for microtubule physiology remain unclear.

Changes in lattice spacing are linked to microtubule physiology in several ways. First, GTP-like lattices are, thus far, frequently expanded, regardless of whether the GTP-like state is maintained by nucleotide analogs (GMPCPP, Paquette et al. 2025 (preprint) 1 (Hyman *et al*. 1995; Alushin *et al*. 2014)) or point mutations that disrupt tubulin’s GTPase activity (LaFrance *et al*. 2022). In contrast, GDP and GDP-Pi lattices are often compacted, suggesting that compaction is coupled to GTP hydrolysis, a central mechanism of microtubule dynamic instability (Carlier *et al*. 1989). An important caveat is that some GTP-like lattices are compacted (Diao *et al*. 2023), while GDP lattices remain expanded in microtubules polymerized from *S. cerevisiae* tubulin (Howes *et al*. 2017), *S. pombe* tubulin (von Loeffelholz *et al*. 2017), and *C. elegans* tubulin (Chaaban *et al*. 2018).

Second, the binding affinity and activity of several microtubule-associated proteins (MAPs) appear to regulate lattice spacing and vice versa. Some MAPs nucleate expanded lattices, bind preferentially to expanded lattices, and/or expand the lattice spacing of pre-formed lattices *in vitro*. For example, kinesin-1 was observed to directly expand pre-formed GDP lattices, creating changes in microtubule length observable by light microscopy (Peet, Burroughs, and Cross 2018), which may impact intracellular trafficking (Shima *et al*. 2018). Other expansion-sensitive MAPs include proteins that regulate templated nucleation and/or minus-end stability (TPX2: (Zhang *et al*. 2017), and CAMSAPs: (Liu and Shima 2023)). Conversely, some MAPs nucleate compacted lattices, bind preferentially to compacted lattices, and/or compact the lattice spacing in conditions where the lattice would otherwise be expanded. These MAPs include the EB-family proteins Bim1 (Howes *et al*. 2017) and Mal3 (von Loeffelholz *et al*. 2017), the neuronal MAP Doublecortin (DCX) (Manka and Moores 2018b), and “cohesive envelopes” of the neuronal MAP tau (Siahaan *et al*. 2022).

Third, lattice spacing regulates the “writers” of post-translational modifications (PTMs) of αβ-tubulin, namely the detyrosinating enzyme VASH (Yue *et al*. 2023) and the acetyltransferase αTAT1 (Shen and Ori-McKenney 2024). Thus, lattice spacing may play a role in the regulation of the tubulin code, which would have downstream impacts on the many MAPs and motor proteins that are regulated by PTMs (Janke and Magiera 2020; Moutin *et al*. 2021).

Finally, changes in lattice spacing can occur when microtubules are bent or curved, for example when microtubules are buckled by compressive forces produced by contractile actin networks (Brangwynne *et al*. 2006). On the concave surface of a bent microtubule, the lattice presumably compresses, producing complex reorganization of tubulin-tubulin contact surfaces (Chew and Cross 2025), while the convex surface of a bent microtubule expands. These reorganizations are recognized, e.g., by proteins that bind preferentially to expanded or compacted lattices (Peet, Burroughs, and Cross 2018).

In these ways, lattice spacing sits at the center of nucleotide hydrolysis, MAP regulation, the tubulin code, and microtubule bending. The picture above suggests that each microtubule, and indeed each αβ-tubulin dimer, is subject to a range of molecular driving forces that alter lattice spacing (e.g., MAP binding, bending forces, etcetera). Some of these driving forces lead to a compacted state (e.g., GTP hydrolysis) and some lead to an expanded state (e.g., kinesin-1 binding). If there is a range of competing forces, how are they reconciled? Are microtubules speckled with dimers in two distinct structural states? Or do microtubules contain separate regions of uniform lattice spacing? The separation of microtubules into distinct local lattice regions has been observed in cells (Tas *et al*. 2017; de Jager *et al*. 2024) and can be accomplished by MAPs such as Tau and MAP7 (Monroy *et al*. 2018), a phenomenon referred to as the “MAP code” (Monroy *et al*. 2020). But the extent to which these distinct MAP-bound regions reflect and/or influence the underlying lattice spacing remains unknown.

In order to study the competition between microtubule expansion and compaction, we needed a model system in which changes in lattice spacing could be reliably induced and studied. Based on previous work, we chose the competition between paclitaxel, a chemotherapeutic drug of the taxane family, and DCX, a neuronal MAP. Paclitaxel nucleates expanded microtubule lattices *in vitro* (Alushin *et al*. 2014), while DCX nucleates compacted lattices (Fourniol *et al*. 2010). In cells expressing DCX-GFP, the addition of paclitaxel caused a significant relocalization of DCX to curved, ring-like segments. This relocalization was interpreted as evidence that paclitaxel caused microtubule expansion, forcing DCX to bind to curved, compacted surfaces (Ettinger *et al*. 2016). This interpretation was subsequently complicated, however, by uncertainty as to whether the addition of paclitaxel can actually cause expansion of pre-formed microtubules (Kellogg *et al*. 2017). Furthermore, paclitaxel and DCX were shown to bind simultaneously to their distinct sites on microtubules *in vitro* (Manka and Moores 2018b); and yet the microtubule lattices were compacted under these conditions. Thus, while it is clear that paclitaxel and DCX have opposing effects on microtubule lattice spacing in isolation, the competitive landscape for these two molecules remains uncharted. We reasoned that charting this landscape could reconcile the disparate findings described above.

In this work, we reconstituted the process of microtubule expansion and compaction *in vitro*, allowing us to directly observe the impact of DCX and paclitaxel on the lattice spacing of pre-formed microtubules by light microscopy. We combined these *in vitro* experiments with a cellular model system in which the localization pattern of DCX serves as a probe for the impact of taxanes on microtubule structure. We find that microtubule lattice spacing depends on the relative concentration of DCX to paclitaxel. When DCX expression is low, high doses of paclitaxel cause a significant relocalization of DCX in response to microtubule expansion, as observed previously (Ettinger *et al*. 2016). However, when DCX concentration is increased sufficiently, microtubule lattices are compacted *in vitro* and in cells, even in the presence of excess paclitaxel. Interestingly, when the relative concentrations of the two molecules are “balanced”, we observed a complex phenotype in which DCX preferentially bound to long, straight microtubules sections that co-existed alongside short bends. These observations suggest that microtubule “expanders” and “compactors” can define local regions of microtubule lattice spacing in this system.

## Results

### Paclitaxel expands microtubules cooperatively, restricting DCX’s localization

To begin our study of microtubule lattice spacing, we developed an *in vitro* buckling assay^1^ to directly observe microtubule expansion using light microscopy. Our assay is similar to pioneering work by Peet *et al*. (2018) and others (Liu and Shima 2023). Briefly, we created “double-capped” microtubules in which a long segment of GDP-lattice is capped at both ends by short segments of stable GMPCPP-lattice anchored to the cover glass (see Methods). We imaged these “double-capped” microtubules by TIRF microscopy. We exchanged the imaging buffer to add 10 µM paclitaxel, and we immediately saw the microtubules buckle and bend outward, indicating that the GDP segment had expanded between the anchored ends (Fig. 1A-B and supplemental movie 1). We measured the microtubule length before and after the addition of paclitaxel. At 10 µM paclitaxel, we measured an increase in microtubule length of *ΔL* = 2.3 % ± 0.1 % (mean ± SEM, *n* = 44). This measurement agrees very well with the lattice spacing changes measured by cryo-EM (2.3 % ± 0.2 %, (LaFrance *et al*. 2022)). In control experiments (buffer exchange only), the GDP segments remained straight, with no observable change in length. We repeated this experiment across a range of paclitaxel concentrations (Fig. 1B-C). Interestingly, as paclitaxel concentration was increased from 0.1 µM to 10 µM, we observed that the length change increased sigmoidally, suggesting a cooperative biochemical process. We fit these *ΔL* measurements to the Hill equation, with a Hill coefficient of *n*_*H*_ = 2.4 ± 0.6. This cooperative behavior is consistent with recent data on a fluorescent taxane, Fchitax-3, which binds to microtubules cooperatively (Rai *et al*. 2020). These data demonstrate that paclitaxel can, in fact, expand microtubules when added after nucleation, in contrast to previous cryo-EM studies (Kellogg *et al*. 2017). Thus, we refer to paclitaxel as a “microtubule expander”.

**Figure 1:**
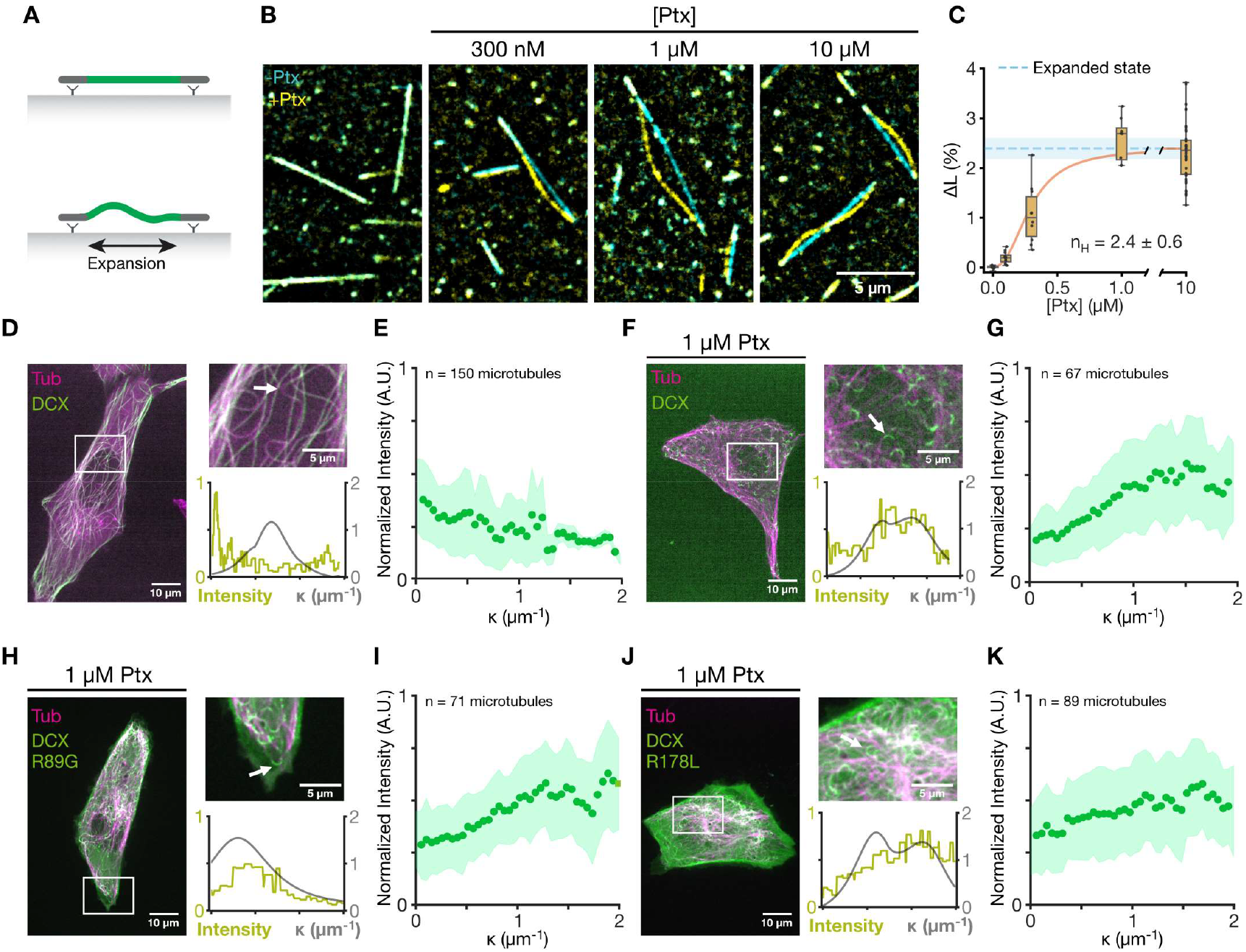
Paclitaxel expands microtubules cooperatively, restricting DCX’s localization. **(A)** Schematic of the microtubule buckling assay. A microtubule containing compacted GDP-tubulin (green) is capped on both ends by TAMRA-labelled GMPCPP-tubulin (grey). GMPCPP-tubulin “caps” are bound to coverslips with anti-TAMRA antibodies. Paclitaxel is flown into the reaction channel triggering the expansion of the GDP-tubulin section. As the GMPCPP caps are fixed to the surface, the microtubule buckles outwards to make up for the increase in length. **(B)** Micrographs of the buckling assay before (cyan) and after (yellow) the addition of paclitaxel (Ptx) at the concentrations indicated. **(C)** Plot of the percentage increase in microtubule length (ΔL) after the addition of paclitaxel as a function of its concentration. Box plot displays median (middle bar), 1^st^ and 3^rd^ quartiles (box limits), and spread of all data (whiskers). Mean and standard error values are fit to a Hill equation. The mean and SEM of cryo-EM measurements of expanded microtubules (LaFrance 2022) are marked by a blue dashed line and blue fill, respectively. n = 15, 12, 10, 8, 44 microtubules pooled from 3 repeats. All mean ΔL per given concentration were significantly different from control, see Table1 for *p* values. **(D), (F), (H), (J)** *Left*: Live cell images of U2OS cells stably expressing α-tubulin-mCherry (magenta) and transfected with DCX-eGFP (green). Cells are untreated (D) or treated with 1 µM paclitaxel (F, H, & J). DCX-eGFP expression is wild type (D & F), R89G mutant (H), or R178L mutant (J). *Right, top*: Insets. White arrows indicate microtubules chosen for representative line scan. *Right, bottom*: Representative line scans of indicated microtubules. DCX intensity (green) and microtubule point curvature κ (grey) are plotted over the length of the trace. **(E), (G), (I), (K)** Normalized fluorescence intensity of DCX-eGFP in U2OS cells plotted as a function of absolute microtubule point curvature κ. Cells are untreated (E) or treated with 1 µM paclitaxel (G, I, & K). DCX-eGFP expression is wild type (E & G), R89G mutant (I), or R178L mutant (K). n = 150 microtubules pooled from 16 cells (E); 67 microtubules pooled from 9 cells (G); 71 microtubules pooled from 8 cells (I); 89 microtubules pooled from 11 cells (K). Datapoints were binned and averaged in intervals of 0.05 μm^-1^. Shaded area represents standard deviation of the mean of each bin.

Next, we used paclitaxel as a microtubule expander to induce changes to microtubule lattice spacing in cells. More precisely, we developed a model system similar to Ettinger *et al*. (2016), where we transfected U2OS cells with a DCX-eGFP construct and added paclitaxel. To start the experiment, we selected cells where DCX-eGFP was readily detectable but did not cause microtubule bundling or other discernible changes to the radial organization of the microtubule array. In these conditions, DCX-eGFP is broadly distributed on microtubules throughout the cell, with a detectable enrichment on straight microtubule segments over curved segments (Fig. 1D-E). Notably, the microtubule network was mostly linear in these DCX-positive cells, while paclitaxel treatment triggered an increase in the distribution of curves (Fig. S1A). When the cells were treated with 1 µM paclitaxel, we observed that DCX-eGFP relocalized to curved, ring-like segments of microtubules (Fig. 1F), consistent with previous observations in cells (Ettinger *et al*. 2016) and *in vitro* (Bechstedt, Lu, and Brouhard 2014) (reproduced in Fig. S1B). We measured the DCX-eGFP intensity as a function of microtubule curvature, κ, using custom image analysis software (Mary and Brouhard 2019) and found that, in the presence of 1 µM paclitaxel, DCX-eGFP intensity increased with increasing κ (Fig. 1G and *in vitro* Fig. S1C). These observations were previously interpreted as evidence that DCX binds preferentially to compacted lattices on the concave surface of the bends. However, DCX also binds preferentially to curved protofilaments at microtubule ends (Bechstedt, Lu, and Brouhard 2014) due to the preference of the DC-2 domain for this region of the lattice (Manka and Moores 2020). To rule out a role for curvature *per se* in these observations, we introduced a point mutation into DC-2 that disrupts DCX’s binding to microtubule ends (R178L, (Bechstedt, Lu, and Brouhard 2014)). We also created a construct with a point mutation in DC-1 as a control (R89G). When we transfected these mutant constructs into U2OS cells and treated the cells with 1 µM paclitaxel, we observed the same phenotypes, namely a relocalization from straight microtubules (Fig. S2A, untreated) to curved, ring-like segments (Fig. 1H-K). Both R89G- and R178L-DCX mutant images and curvature-intensity correlations were similar to wildtype DCX data. This result indicates that the relocalization of DCX induced by paclitaxel does not depend on DC-2’s preference for protofilament curvature found at microtubule ends (Bechstedt, Lu, and Brouhard 2014). Rather, we concur with the interpretation that DCX binds to the curved segments because of compaction on their concave surface. As high concentrations of paclitaxel compete for lattice spacing, most lattices become expanded; DCX relocalizes to the only remaining compacted regions, which are found at curved regions of the lattice (Ettinger *et al*. 2016; Peet, Burroughs, and Cross 2018).

### Lattice expansion by stabilizing drugs relocalizes DCX

Before continuing our investigation of the competition between DCX and paclitaxel, we paused to ask whether *any* molecule that binds the taxane pocket would expand the microtubule lattice as paclitaxel does. To this end, we tested three additional drugs that bind to the taxane site for their ability to (a) expand GDP-microtubules *in vitro* and (b) relocalize DCX-eGFP in cells. Overall, whenever we observed expansion of GDP-microtubules *in vitro*, we also observed DCX-eGFP relocalize to curved microtubule segments in cells, confirming that DCX responds to local lattice spacing.

First we tested docetaxel, a member of the taxane family of tetracyclic diterpenoids, which differs only slightly from paclitaxel in terms of the side chains of the taxane ring (Fig. S3A-B, blue). Upon addition of docetaxel to the *in vitro* buckling assay, the GDP-microtubule length increased (Fig. 2A). *ΔL* = 2.5 % ± 0.1 % was reached at 0.3 µM docetaxel (mean ± SEM, n = 28). Thus, docetaxel is also a microtubule expander. Relative to paclitaxel, docetaxel’s effect plateaued at lower concentrations, consistent with docetaxel’s 3-fold higher affinity for the taxane site (Buey *et al*. 2005). When 1 µM docetaxel was added to U2OS cells expressing DCX-eGFP, we observed DCX relocalize to curved microtubule segments (Fig. 2B-C, *in vitro* Fig. S1D-E), indicating that the correlation between *in vitro* expansion and relocalization of DCX in cells is not unique to paclitaxel.

**Figure 2:**
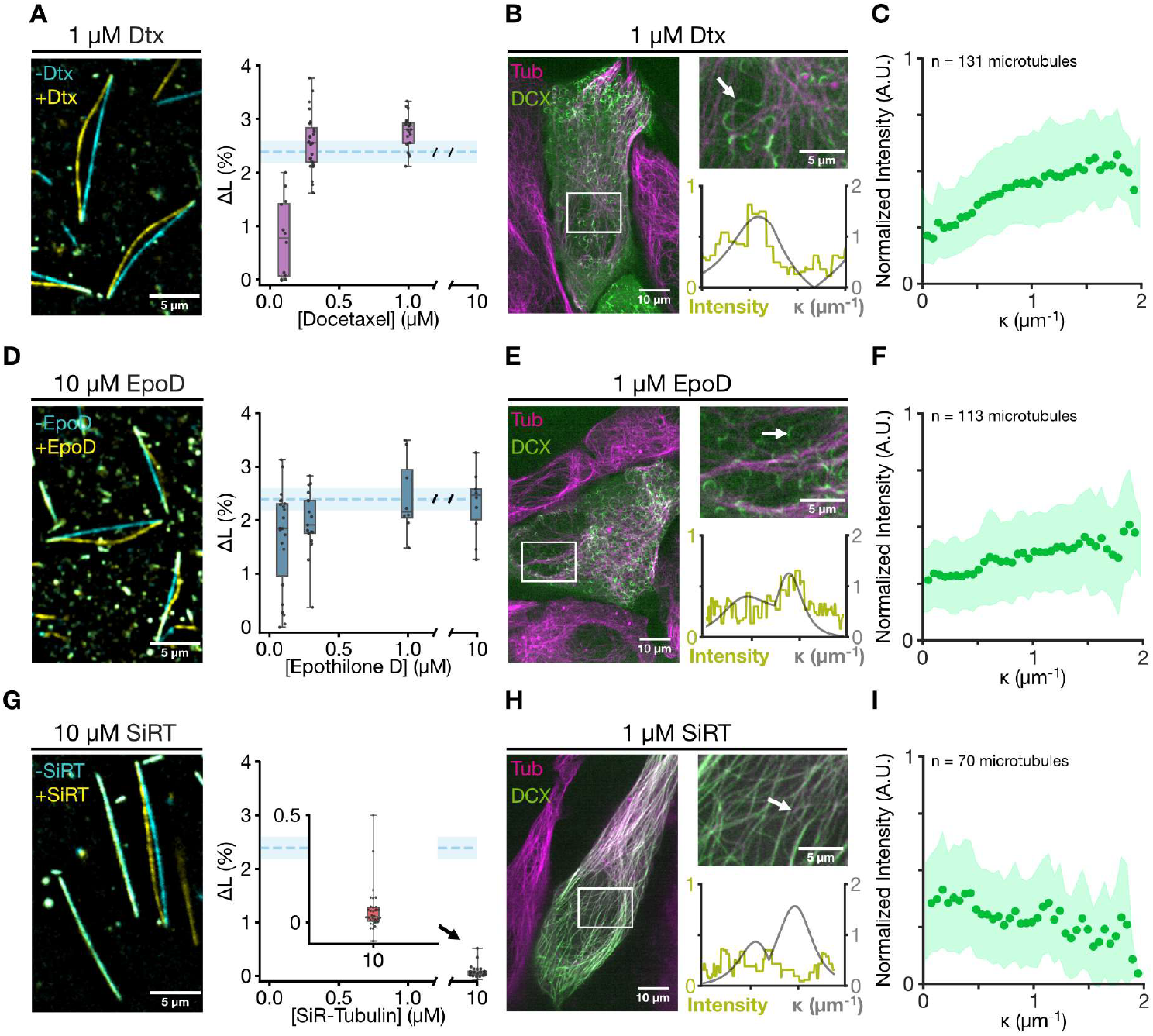
Lattice expansion by stabilizing drugs relocalizes DCX. **(A), (D), (G)** *Left*: Micrographs of GDP-tubulin microtubules before (cyan) and after (yellow) the addition of 10 µM docetaxel (Dtx) (A), 10 µM epothilone D (EpoD) (D), or 10 µM SiR-tubulin (SiRT) (G). *Right*: Plots of the percentage increase in microtubule length (ΔL) after the addition of each drug as a function of drug concentration. Blue dashed lines and fill mark the expansion percentage as in Figure 1C. n = 12, 28, 20 microtubules pooled from 3 repeats (A); 22, 17, 8, 10 microtubules pooled from 3 repeats (D); 32 microtubules pooled from 3 repeats (G). All mean ΔL per given concentration were significantly different from control, see Table1 for *p* values. **(B), (E), (H)** *Left*: Live cell images of U2OS cells stably expressing α-tubulin-mCherry (magenta), transfected with DCX-eGFP (green) and treated with 1 µM docetaxel (B), 1 µM epothilone D (E), or 1 µM SiR-tubulin (H). *Right, top*: Insets. White arrows indicate microtubules chosen for representative line scan. *Right, bottom*: Representative line scans of indicated microtubules. DCX intensity (green) and microtubule point curvature κ (grey) are plotted over the length of the trace. **(C), (F), (I)** Normalized fluorescence intensity of DCX-eGFP in U2OS cells plotted as a function of absolute microtubule point curvature κ. Cells are treated with 1 µM docetaxel (C), 1 µM epothilone D (F), or 1 µM SiR-tubulin (I). n =131 microtubules pooled from 15 cells (C); 113 microtubules gathered from 12 cells (F); 70 microtubules gathered from 9 cells (I). Datapoints were binned and averaged in intervals of 0.05 μm^-1^. Shaded area represents standard deviation of the mean of each bin.

Next we tested epothilone D, a member of the epothilone family of 16-membered macrolides, which binds the taxane pocket (Giannakakou *et al*. 2000; Bollag *et al*. 1995) (Fig. S3C). In the *in vitro* buckling assay, epothilone D caused GDP-microtubules to expand. *ΔL* = 2.4 % ± 0.3 % was reached at 1 µM epothilone D (mean ± SEM, n = 17) (Fig. 2D). As with docetaxel, epothilone-D expanded microtubules at lower concentrations than paclitaxel, suggesting a higher affinity. In cells, addition of 1 µM epothilone-D caused DCX-eGFP to relocalize (Fig. 2E-F, *in vitro* Fig. S1F-G). Thus, the correlation between *in vitro* expansion and re-localization of DCX in cells extends to three molecules that bind the taxane site.

We obtained different results, however, with SiR-tubulin, a derivative of docetaxel in which a Si-Rhodamine fluorophore is linked to the C13 side chain of the taxane ring (Lukinavičius *et al*. 2016) (Fig. S3D, blue). Interestingly, even when 10 µM SiR-tubulin was added to the *in vitro* buckling assay, only limited changes in length were observed (Fig. 2G, *ΔL* = 0.06 % ± 0.02 % (mean ± SEM, n = 32)). Consistently, when we added 1 µM SiR-tubulin to U2OS cells expressing DCX-eGFP, the DCX signal remained on straight microtubules throughout the cell (Fig. 2H-I, *in vitro* Fig. S1H-I). Thus, we conclude that SiR-tubulin is not a microtubule expander in our experiments, a result we discuss further below.

### Paclitaxel and DCX compete for control of lattice spacing

Thus far, our results demonstrate clearly that paclitaxel and other drugs are microtubule expanders that, when used at high concentrations, relocalize DCX to curved, compacted regions of the lattice. However, a recent cryo-EM study showed that microtubules copolymerized with 3.5 µM DCX and subsequently stabilized with 1 mM paclitaxel maintained a compacted lattice spacing (Manka and Moores 2018b), indicating that DCX can dominate lattice spacing at sufficiently high concentrations *in vitro*. We wondered if we could find additional evidence to support this observation.

To investigate this idea, we introduced purified recombinant DCX into the *in vitro* buckling assay. First, we expanded GDP-microtubules by adding an excess of paclitaxel (10 µM, *ΔL* = 2.3 ± 0.1 % (mean ± SEM, *n* = 44)). Then, we exchanged the buffer to include both 10 µM paclitaxel and 500 nM recombinant DCX-eGFP. Upon exchange, the microtubules returned to their original length measured at the start of the experiment (*ΔL* = 0.1 % ± 0.1 % (mean ± SEM, n = 12)) (Fig. 3A-B). From this result, we conclude that a high concentration of DCX can compact a GDP-microtubule expanded by paclitaxel *in vitro*, even in presence of excess paclitaxel.

**Figure 3:**
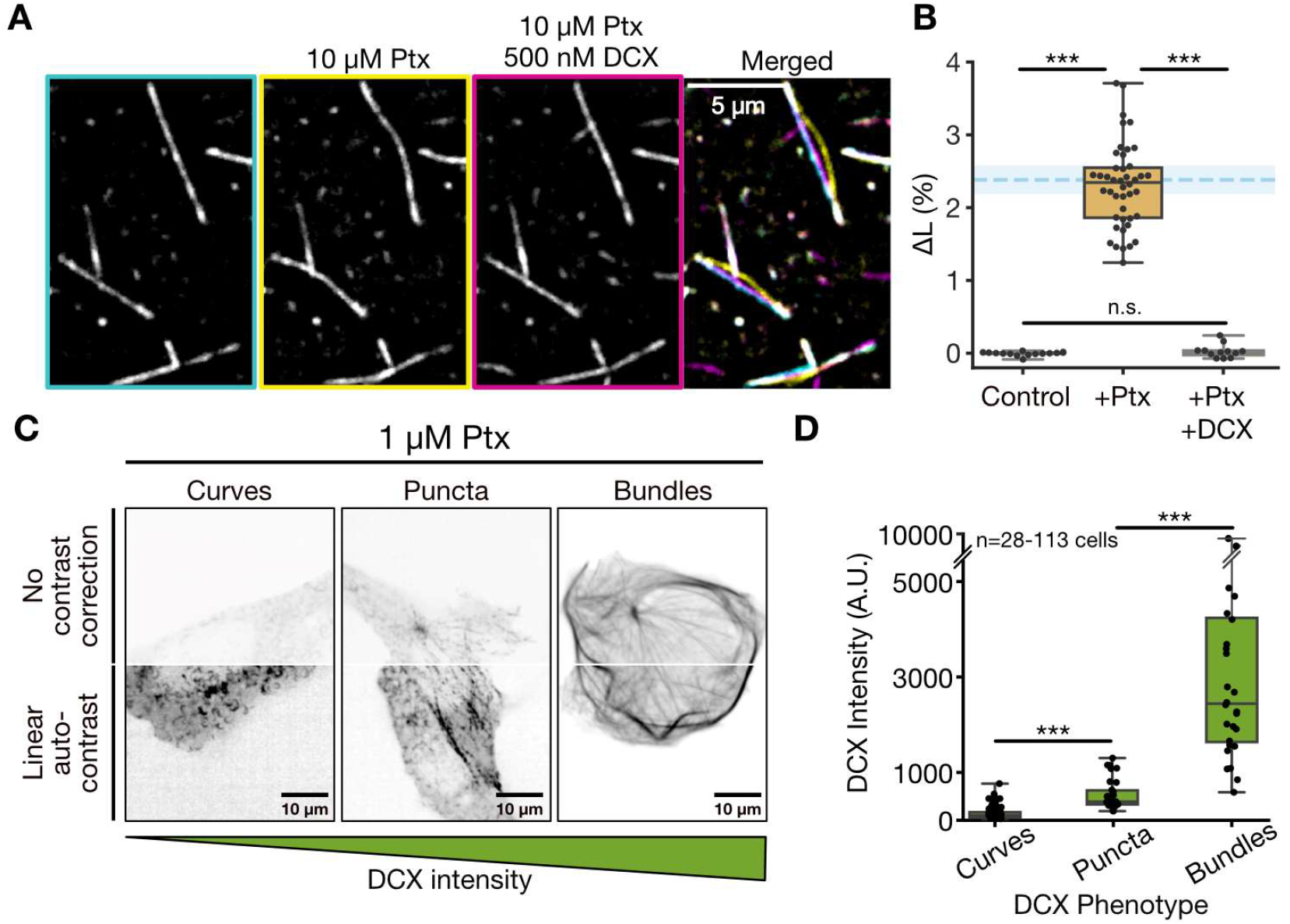
Paclitaxel and DCX compete for control of lattice spacing. **(A)** Micrographs of GDP-tubulin microtubules before treatment (cyan), after the addition of 10 µM paclitaxel (yellow), and after the addition of 10 µM paclitaxel and 500nM DCX (magenta). **(B)** Plot of the percentage increase in microtubule length (ΔL) after the addition of paclitaxel or paclitaxel and DCX. Blue dashed line and fill mark the expansion percentage as in Figure 1C. n = 15, 44, 12 microtubules pooled from 3 repeats. Statistical analysis was done using Welch’s one-sided t tests, n.s. nonsignificant, *** *p* < 0.001. See Table 1 for *p* values. **(C)** Representative images of DCX-eGFP in U2OS cells stably expressing α-tubulin-mCherry, transfected with DCX-eGFP and treated with 1 µM paclitaxel. *Top row*: Unprocessed images. *Bottom row*: Images processed with linear auto-contrast to enhance observation of the DCX phenotype. **(D)** Mean DCX fluorescence intensity in each DCX phenotype category detailed in C. Box plot displays median (middle bar), 1^st^ and 3^rd^ quartiles (box limits), and spread of all data (whiskers). n = 113, 30, and 28 cells. Statistical analysis was done using Welch’s one-sided t tests: *** *p* < 0.001.

We wondered if we could observe similar dominance of DCX over paclitaxel in cells. In our previous experiments in U2OS cells described above, we had selected only relatively “low-expressing” cells free from any observable microtubule bundling. To examine higher levels of DCX expression, we optimized our imaging protocols to assess a broad range of DCX-eGFP levels across a population of cells (see Methods). Figure 3C shows several cells at constant paclitaxel concentration (1 µM) but at different DCX-eGFP expression levels. The top half of the image shows the unprocessed image data (to display unmodified signal intensity), while the bottom half is contrast-corrected (to display DCX-eGFP localization). Interestingly, multiple DCX localization patterns were visible, and each pattern corresponded to a specific range of DCX-eGFP expression (Fig. 3C). To quantify this observation, we assigned a category to the localization pattern of DCX and then measured the intensity of DCX-eGFP within each category (Fig. 3D). At low DCX expression levels, 1 µM paclitaxel caused DCX to relocalize to curved, ring-like segments, as described above (Fig. 3C-D, Curves). However, at medium DCX expression levels, the addition of 1 µM paclitaxel caused DCX-eGFP to relocalize to bright puncta and clusters, which sometimes formed long, straight assemblies along the microtubules, but which did not cover the whole microtubule network (Fig. 3C-D, Puncta). At high DCX expression levels, DCX-eGFP formed microtubule bundles and appeared to cover the entire microtubule network (Fig. 3C-D, Bundles). Importantly, this “high-expression” phenotype did not change with the addition of 1 µM paclitaxel (Fig. 3C-D, Bundles, compare to Fig. S2B, DCX overexpression with no paclitaxel). Thus, high levels of DCX expression appear to resist relocalization and the formation of curved microtubule segments. Combining these data with our *in vitro* results (Fig. 3A-B), we conclude that DCX can resist paclitaxel-induced expansion when it is present at sufficient levels.

**Table 1:** Mean and SEM values of ΔL. Statistical analysis was done using Welch’s one-tailed t test.

| Drug | [Drug] (μM) | [DCX] (nM) | Mean ΔL | SEM ΔL | n | p-value compared to control |
| --- | --- | --- | --- | --- | --- | --- |
| Control | 0 | 0 | -0.00566 | 0.007015 | 15 |  |
| Paclitaxel | 0.1 | 0 | 0.181421 | 0.033646 | 12 | 7.55E-05 |
| Paclitaxel | 0.3 | 0 | 1.052885 | 0.186107 | 10 | 1.49E-04 |
| Paclitaxel | 1 | 0 | 2.571357 | 0.155771 | 8 | 3.48E-07 |
| Paclitaxel | 10 | 0 | 2.300036 | 0.086749 | 44 | 8.52E-29 |
| Docetaxel | 0.1 | 0 | 0.774085 | 0.214739 | 12 | 1.97E-03 |
| Docetaxel | 0.3 | 0 | 2.554492 | 0.097005 | 28 | 3.13E-21 |
| Docetaxel | 1 | 0 | 2.761353 | 0.07196 | 20 | 5.15E-20 |
| Epothilone D | 0.1 | 0 | 1.658166 | 0.202788 | 22 | 2.72E-08 |
| Epothilone D | 0.3 | 0 | 1.96837 | 0.147349 | 17 | 1.97E-10 |
| Epothilone D | 1 | 0 | 2.439872 | 0.255322 | 8 | 1.41E-05 |
| Epothilone D | 10 | 0 | 2.305376 | 0.192127 | 10 | 3.70E-07 |
| SiR-tubulin | 10 | 0 | 0.057693 | 0.019081 | 32 | 1.73E-03 |
| Paclitaxel | 10 | 500 | 0.027292 | 0.027055 | 12 | 1.30E-01 |

### Balanced ratios of DCX and paclitaxel create complex local regions of DCX

The results above suggest that the change in DCX’s localization pattern in response to taxanes depends on the relative concentrations of DCX and paclitaxel. When DCX expression is low, high doses of paclitaxel cause a significant relocalization of DCX in response to microtubule expansion, as observed previously (Ettinger *et al*. 2016) and in a different cell type (RPE-1 cells, Fig. S4A-B). However, when DCX concentrations are increased sufficiently, microtubule lattices are compacted *in vitro* and in cells, even in the presence of excess paclitaxel. We were curious about DCX’s localization pattern when the relative concentrations are more “balanced”: where DCX and paclitaxel are both present, but neither species has overwhelmed the microtubule cytoskeleton, e.g. by freezing dynamic instability (paclitaxel) or by forming microtubule bundles (DCX).

To determine the localization pattern of DCX under “balanced” conditions, we treated low DCX-expressing cells with 90 nM paclitaxel, as shown in Figure 4A, a concentration at which dynamic instability is still observed. We observed a complex and multi-faceted phenotype in which DCX localized to multiple different local regions (Gallery Fig. S2C) independently of the cell type used (Fig. S4C). The brightest signal came from a handful of long, straight microtubule segments that were observed (a) clustered near the cell periphery or (b) growing through the cytoplasm (Fig. 4A, unfilled triangle). These microtubules were reminiscent of the “light sabers” of DCX-eGFP observed on growing microtubules *in vitro* (Bechstedt, Lu, and Brouhard 2014).

**Figure 4:**
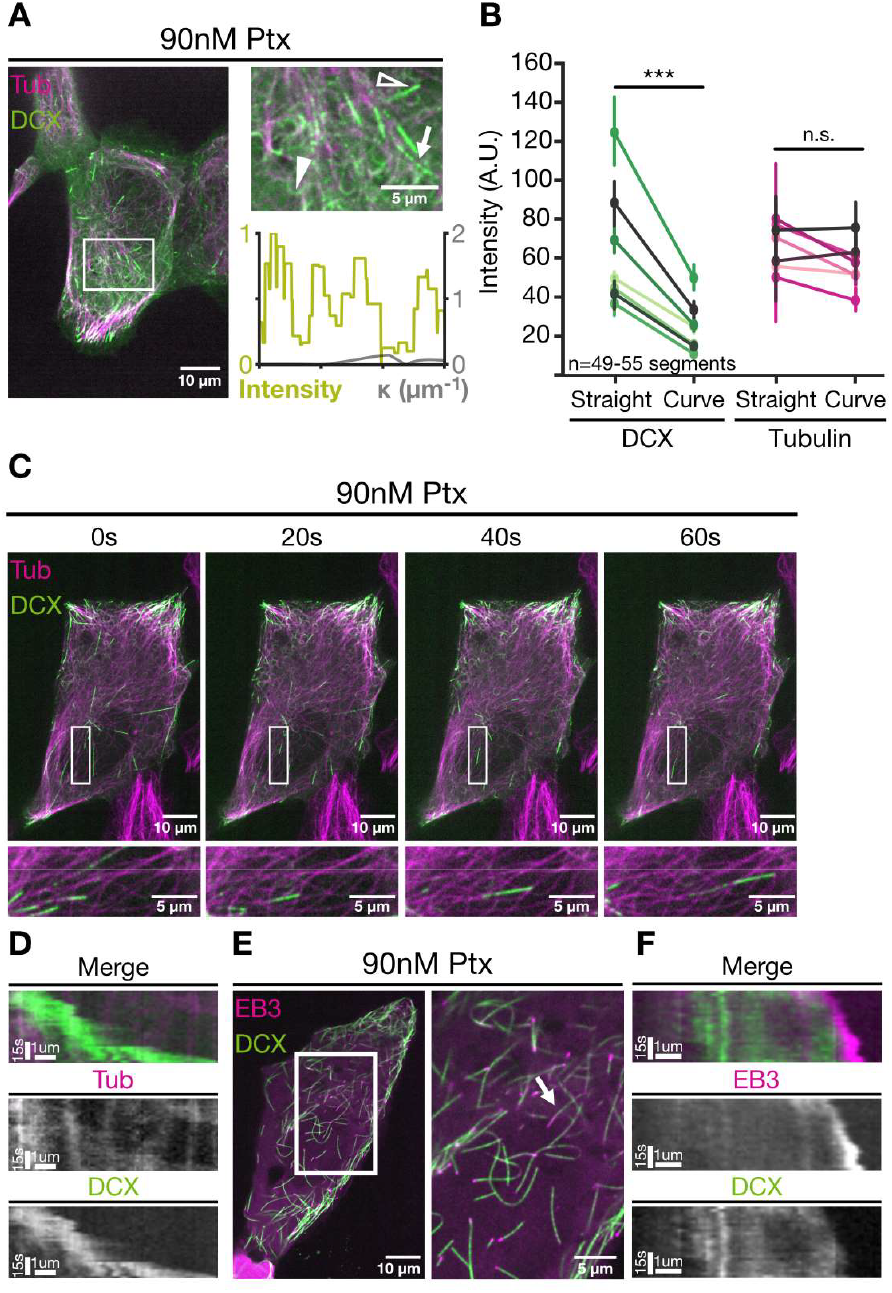
Expanded and compacted lattices co-exist in cells. **(A)** *Left*: Live cell image of U2OS cells stably expressing α-tubulin-mCherry (magenta), transfected with DCX-eGFP (green) and treated with 90 nM paclitaxel. *Right, top*: Inset. Filled triangle indicates ring-like microtubule segment. Unfilled triangle indicates straight microtubule segment. White arrow indicates microtubule chosen for representative line scan. *Right, bottom*: Representative line scan of indicated microtubule. DCX intensity (green) and microtubule point curvature κ (grey) are plotted over the length of the trace. **(B)** Mean intensity of DCX-eGFP and α-tubulin-mCherry at straight and at curved microtubule segments in U2OS cells displaying both phenotypes after treatment with 90 nM paclitaxel. Intensities from the same cell are paired using a line. n = 104 segments from 7 cells. Statistical analysis was done using Wilcoxon signed-rank test: n.s. nonsignificant, *** *p* < 0.001. **(C)** Live imaging time series of a U2OS cell stably expressing α-tubulin-mCherry (magenta), transfected with DCX-eGFP (green) and treated with 90 nM paclitaxel. White boxes show the growing microtubule presented below D. **(D)** Kymograph generated from the microtubule highlighted in C. DCX covers the growing end of the microtubule. **(E)** Live cell image and inset (white frame) of a U2OS cell transfected with EB3-mRuby (magenta) and DCX-eGFP (green) and treated with 90 nM paclitaxel. White arrow indicates the microtubule chosen for representative kymograph in F. **(F)** Kymograph generated from microtubule highlighted in F. EB3 is distal from DCX in growing microtubules.

Although not as bright, we also observed that DCX localized to curved, ring-like segments similar to those observed previously (Fig. 4A, filled triangle). More precisely, we found that DCX was approximately half as bright at curved segments relative to the straight extensions (Fig. 4B, DCX, straight: 63.8 A.U. ± 8.88 A.U. vs curves: 28.7 A.U. ± 4.40 A.U., *p*< 0.01, one-sided Wilcoxon signed-rank test), while tubulin intensity did not vary between these locations (Fig. 4B, tubulin). We posit this 2-fold difference results from DCX covering the whole circumference of the lattice at straight extensions, but only half of it at ring-like curves, since the concave side is compacted and the convex side expanded (Ettinger *et al*. 2016; Peet, Burroughs, and Cross 2018). Finally, many microtubule sections had no detectable DCX signal at all, suggesting that paclitaxel had successfully expanded them.

We looked more closely at the dynamics of the long, bright DCX extensions observed on growing microtubules (See supplemental movie 2). We noticed, for example, that the length of the extensions varied over time, sometimes increasing in length and sometimes shrinking (Fig. 4C-D). Over tens of seconds, the bright extensions would sometimes fragment into immobile clusters (Fig. 4C-D); sometimes they disappeared completely (see supplemental movie 2). These observations indicate that DCX localization is dynamic in the presence of 90 nM paclitaxel, most likely due to rapid changes in the affinity of DCX for local regions of the microtubule cytoskeleton. We wondered if this phenomenon was akin to the “tip-tracking” behavior observed by +TIP proteins (Akhmanova and Steinmetz 2008) and by DCX *in vitro* (Bechstedt and Brouhard 2012). We co-transfected U2OS cells with an EB3-mRuby construct and DCX-eGFP. When we added 90 nM paclitaxel to these cells, we observed EB3-mRuby “comets” at the ends of microtubules with bright DCX extensions (Fig. 4E-F). In kymographs of individual microtubules, the DCX-eGFP signal trails behind the EB3-mRuby comet. We conclude that EB3-mRuby continues to decorate the “GTP cap” in the presence of 90 nM paclitaxel, and that DCX preferentially binds to growing microtubules just behind the region decorated by EB proteins (Ettinger *et al*. 2016).

Thus, at balanced ratios of DCX and paclitaxel, the cells appear to contain separate regions of uniform lattice spacing. In other words, lattice spacings are not randomly speckled but rather organized into distinct local regions of the microtubule across multiple experimental contexts.

## Discussion

In these experiments, we established *in vitro* and cellular model systems for exploring a competition for control of microtubule lattice spacing between a microtubule “expander” (paclitaxel) and a microtubule “compactor” (DCX). We discovered a complex competitive landscape, in which DCX’s localization pattern in cells varied significantly depending on the relative concentration of DCX to paclitaxel (Fig. 3). Importantly, at “balanced” concentrations, DCX preferentially localized to a subset of long, straight microtubules that co-existed alongside microtubules without detectable DCX signal (Fig. 4). We interpret these findings in light of *in vitro* reconstitution experiments in which we could directly observe paclitaxel and DCX cause changes in microtubule lattice spacing (Fig. 1, 3). Ultimately, DCX is the stronger of the two molecules, in that high concentrations of DCX will compact microtubules even in the presence of high concentrations of paclitaxel (Fig. 3) (Manka and Moores 2018b).

From a synthetic biology perspective, our results are a proof of concept for the coexistence of microtubules with distinct lattice spacing in cells (Fig. 4). Our findings corroborate recent cryo-electron tomography results demonstrating the coexistence of distinct lattices using the stableMARK reporter (de Jager *et al*. 2024; Jansen *et al*. 2023). These phenotypes are similar to the separate regions of lattice binding formed by MAPs such as tau and MAP4, or DCX and MAP7 (Siahaan *et al*. 2022; Monroy *et al*. 2018). More broadly, our work contributes to a growing body of evidence documenting the side-by-side coexistence of microtubules with distinct properties (Tas *et al*. 2017; Vinopal *et al*. 2023; Alvarez Viar *et al*. 2024; de Jager *et al*. 2024), where the properties include a microtubule’s complement of MAPs, PTMs, and lattice spacings. Microtubule coexistence is not trivial to explain, because it runs counter to the expectation of randomness arising from diffusion and stochastic reaction pathways. Such randomness should produce microtubules that are stochastically speckled with MAPs, PTMs, and lattice spacings, not distinct local regions. Rather, local regions of, e.g., uniform lattice spacing require a positive feedback mechanism wherein a distinct region of the microtubule reinforces its identity. A simple positive feedback mechanism is cooperative binding. DCX binds cooperatively to microtubules (Bechstedt and Brouhard 2012; Rafiei *et al*. 2022), and microtubule expansion by paclitaxel is cooperative as well (Fig. 1), consistent with data on Fchitax-3 (Rai *et al*. 2020). It will be interesting to know if other examples of microtubule coexistence are explained by cooperative binding mechanisms.

Our results may also help explain the mechanism of action of taxanes in clinically-relevant contexts. Taxanes both expand the microtubule lattice (Fig. 1, (Alushin *et al*. 2014)) and stabilize the “M-loops” that comprise the lateral bonds (Prota *et al*. 2023). Clinically, taxanes are widely assumed to function as inhibitors of microtubule dynamic instability and mitotic progression. This assumption is at odds with their impact on solid tumors with a low mitotic index, a conundrum termed the “proliferation rate paradox” (Mitchison 2012). This paradox remains unsolved, in part because the concentration of paclitaxel in patient blood plasma is sub-µM, much lower than the concentrations used in some tissue culture experiments. In our experiments, low concentrations of paclitaxel produced significant changes to DCX localization. We speculate that other MAPs may be similarly relocalized when cells are treated with taxanes, which could disrupt cell physiology in complex ways.

Two other taxanes behaved as microtubule expanders in our assays, namely docetaxel and epothilone D, but the docetaxel-derivative SiR-tubulin did not (Fig. 2). The SiR-tubulin results are encouraging, considering that SiR-tubulin is marketed as a “non-disruptive” fluorescent marker for microtubule dynamics in cells. However, previous *in vitro* experiments have suggested that SiR-tubulin’s affinity for microtubules varies depending on whether MAPs are bound. For example, SiR-tubulin signal is lower on microtubules with tau or MAP2c (Siahaan *et al*. 2022) and higher on microtubules with MAP7 or αTAT1 (Shen and Ori-McKenney 2024). This variation in SiR-tubulin’s affinity points towards a preferential binding to expanded microtubule lattices. That said, in our experiments, we observed colocalization of SiR-tubulin signal and DCX signal, both *in vitro* and in cells. We conclude that fluorescent taxanes remain powerful, if complex, tools for studying microtubules *in vitro* and in cells. An important limitation of our study is its synthetic nature. DCX and paclitaxel are not natural competitors: DCX is normally expressed only in developing neurons (Gleeson *et al*. 1999; Francis *et al*. 1999) and paclitaxel does not cross the blood-brain barrier, as it is exported from brain endothelial cells by P-glycoprotein (Fellner *et al*. 2002; Woo *et al*. 2003). But DCX remains a sensitive probe for changes in microtubule structure, as demonstrated here for lattice spacing and in previous work for curvature at microtubule ends (Bechstedt, Lu, and Brouhard 2014) and nucleotide state (Manka and Moores 2020).

At present, relatively few MAPs have been tested for their sensitivity to microtubule lattice spacing relative to the large microtubule interactome. But the initial catalog includes such critical proteins as kinesin-1, tau, MAP2c, TPX2, CAMSAP3, VASH, αTAT1, and of course DCX. Notably, the initial catalog includes a significant diversity of protein domains and corresponding contact surfaces on the microtubule lattice (Manka and Moores 2018a). In other words, sensitivity to lattice spacing is found across protein families, suggesting widespread adaptation of protein domains to this structural feature of microtubules. Recently, we showed that DCX, a microtubule compactor, regulates α-tubulin polyglutamylation in neurons (Sébastien *et al*. 2025). We speculate that MAPs able to control lattice spacing in cells participate in defining tubulin PTMs (Shen and Ori-McKenney 2024; Yue *et al*. 2023) and microtubule patterns (van Grinsven and Akhmanova 2025). We look forward to exploring the complete microtubule interactome in this new light.

## Acknowledgements

We would like to thank current members of the Brouhard, Hendricks, and Bechstedt labs, for fruitful discussions during all phases of this project. We would also like to acknowledge the McGill University Advanced BioImaging Facility (ABIF), RRID: SCR_017697 and the Genome Québec sequencing platform and their personnel for providing services and helpful advice for data collection and analysis. G.J.B. acknowledges support from Natural Sciences and Engineering Research Council of Canada (RGPIN-2014-03791), the Canadian Institutes of Health Research (PJT-148702), and McGill University. A.G.H. is supported by the Canadian Institutes of Health Research (PJT-185997). This paper was typeset with the bioRxiv word template by @Chrelli:www.github.com/chrelli/bioRxiv-word-template

## Author contributions

Conceptualization and experimental design: G.J.B., M.S., A.L.P, and S.C.T. Experimental procedures: A.L.P., S.C.T., and J.A.G.H. Code writing: S.C.T. and A.L.P. Data analysis: A.L.P. and S.C.T. Figures and schematics design: A.L.P. and S.C.T. Manuscript writing: A.L.P, S.C.T, M.S., and G.J.B. All authors participated in editing the manuscript. Funding acquisition: G.J.B. and A.G.H.

## Materials and Methods

### Cell culture and transfection

Cellular assays were performed in human osteosarcoma cells (U2OS) and human retinal pigment epithelial cells (RPE-1) maintained at 37°C/5% CO2 in DMEM-high glucose media (Gibco, #11995-065) supplemented with 10% fetal bovine serum (Sigma-Aldrich, #F1051), and 1% penicillin/streptomycin (Wisent, #450-201-EL). The U2OS cell line stably expressing BAC-based α-tubulin-mCherry under a CMV promoter (gift from Alex Bird) was selected with G418. Cells were passaged with 0.05% trypsin (Gibco, #25300-054) and seeded in glass-bottom dishes or 96-well plates.

The calcium phosphate method (Graham and van der Eb 1973) was used to express DCX-eGFP and EB3-mRuby under a CMV promoter in cells immediately after seeding. Constructs were transfected at 3 µg DNA/glass-bottom dish, or 0.3µg DNA/96-well. 24 hours after transfection, cells were rinsed twice with PBS and supplemented with fresh media.

### Live cell taxane assay

Transfected cells (above) were treated with either 1 µM or 90 nM paclitaxel (European Pharmacopoeia, #Y0000698), 1 µM docetaxel (Abcam, #ab141248), 1 µM epothilone D (Abcam, #ab143616), or 1 µM SiR-tubulin supplemented with 10 µM verapamil (Spirochrome, #SC002). Drugs were stored in DMSO and diluted in fresh media to the indicated concentration. Drugs were applied to the cells for 20-30 minutes (paclitaxel, docetaxel, epothilone D) or 45-60 minutes (SiR-tubulin) before imaging and were left in for the duration of imaging (15-30 minutes).

### Live cell imaging

Microscopes used for cellular imaging in this manuscript are part of the McGill University Advanced BioImaging Facility (ABIF), RRID:SCR_017697. Live single cell images and movies were acquired with a Leica DMI6000B inverted microscope equipped with a Quorum WaveFX-X1 confocal spinning disk system on a Photometrics PRIME BSI CMOS camera using a 63x/1.4 Plan-Apochromat oil objective. Cells were kept at 37°C/5% CO_2_ during imaging using a Live Cell Instruments Chamlide TC environmental control system. Excitation wavelengths were λ=491nm (GFP), λ=568nm (mCherry and mRuby), and λ=643nm (Si-rhodamine). Images and movies were acquired with Metamorph software on a single focal plane at the bottom of the cell where microtubules below the nucleus were in focus. Channels were acquired at 500ms exposure (DCX-eGFP, Tub-mCherry) or 400ms exposure (EB3-mRuby). Exposure times and laser power were kept constant between imaging sessions and conditions. Movie frames were acquired every two seconds for 60-120 seconds. Cells selected for imaging were selected such that individual microtubules could be observed and remained unsaturated throughout the acquisition.

High content screening live cell images were acquired with a Perkin-Elmer OperaPhenix confocal spinning disk system on sCMOS cameras using a 20X NA 1.0 water objective. Cells were kept at 37°C/5% CO_2_ during imaging using the OperaPhenix environmental control system. Excitation wavelengths were λ=488nm (GFP) and λ=561nm (mCherry). Images were acquired with Harmony 9.0 software on a single focal plane at the bottom of the cell where microtubules below the nucleus were in focus. No bias was used in selecting imaging areas.

### Protein purification

Tubulin was purified from juvenile bovine brains by cycles of polymerization and depolymerization followed by loading onto a column containing Fractogel EMD SO ^-^ (M) resin (Millipore Sigma), as described previously (Chaaban *et al*. 2018). Tubulin was either cycled once more and aliquoted to make unlabelled tubulin, or cycled twice with an intermediate incubation with fluorescent dyes (tetramethylrhodamine (TAMRA), Atto 633 or Atto 550 succinimidyl esters) to make labelled tubulin as described by Hyman *et al*. (1991).

Recombinant His-DCX-eGFP-Strep (accession number NP_835365, Addgene #83918) was expressed in BL21(DE3) *E. coli* (New England Biolabs) at OD(600) = [0.4-0.6] and induced with 0.4 mM IPTG at 18 °C overnight. Cells were harvested by centrifugation, resuspended in Buffer A [50 mM NaPO_4_, 5 mM Imidazole, 300 mM NaCl, 5% Glycerol (v/v), pH 8.0] and lysed in an EmulsiFlex-C5 (Avestin). Cleared lysate was loaded onto a His60 Ni Superflow Resin column (Takara Bio) and eluted using Buffer A with a final concentration of 200 mM Imidazole and 10% glycerol (v/v). His-elution was subsequently diluted 1:5 into Buffer W [100 mM Tris-HCl, 150 mM NaCl, 1 mM EDTA, pH 8.0], loaded into Strep-Tactin Sepharose (IBA Lifesciences) and eluted with Buffer W supplemented with 2.5 mM D-Desthiobiotin and 10% Glycerol (v/v). Protein was loaded on a size exclusion chromatography column (Cytiva Superdex 200 Increase 10/300 GL). The peak fraction with the smallest mass (p) was recovered and concentrated. Protein aliquots were flash frozen in liquid nitrogen. Concentration was determined by absorbance at 280 nm with a DS-11 FX spectrophotometer (DeNovix) and purity was determined by SDS-PAGE (Fig. S5).

### Microtubule reconstitution assay

Microtubules were visualized bound to the surface of a glass coverslip as described previously (Gell *et al*. 2010). Coverslips were cleaned, silanized, and used to make solution exchange channels on custom-made microscope holders as described previously (Chaaban *et al*. 2018; Helenius *et al*. 2006). GMPCPP microtubule seeds were prepared by polymerizing a 1:15 molar ratio of TAMRA labelled:unlabelled tubulin in the presence of guanosine-5’-[(α,β)-methyleno]triphosphate (GMPCPP, Jena Biosciences) in two cycles, as described (Gell *et al*. 2010). Drug-stabilized microtubules were prepared by polymerizing a 1:20 molar ratio of Atto 633 or Atto 550 labelled:unlabelled tubulin, then diluting the reaction 1:32 into BRB80 (80 mM PIPES-KOH pH 6.9, 1 mM EGTA, 1 mM MgCl_2_) supplemented with 10 µM of drug, pelleted and resuspended in a suitable volume of 10 µM drug-BRB80.

Channels were prepared by incubating one channel volume of anti-TRITC (ThermoFisher Scientific, #A6397) diluted 1:25 to bind GMPCPP seeds or anti-β-tubulin antibody (Sigma-Aldrich, #T4026) or diluted 1:10 to bind drug-stabilized microtubules, for 5 minutes. Then the channel was blocked with 1% Pluronic F-127 for 20 minutes and finally, GMPCPP-seeds or drug-stabilized microtubules were incorporated by flowing in one channel volume and incubating for 2-5 minutes. The temperature of the channel was kept at 34 °C, measured using a thermocouple (Omega). On each day of experiments, large aliquots of tubulin and DCX were thawed, sub-aliquoted, and stored in liquid nitrogen. One sub-aliquot of each protein was thawed for each experiment.

Double capped microtubules were made by growing dynamic microtubules from GMPCPP seeds. Flow channels were incubated with 10 µM tubulin (10% Atto 633 or 550 labelled) in imaging buffer (BRB80, 0.1 mg/mL BSA, 10 mM dithiothreitol, 0.01% methylcellulose, 250 nM glucose oxidase, 64 nM catalase, and 40 mM D-glucose) supplemented with 1 mM GTP. After 15 minutes, buffer was exchanged with pre-warmed GMPCPP buffer (0.5 µM tubulin 10% TAMRA labelled, in imaging buffer supplemented with 1 mM GMPCPP). The channel was incubated for 5 minutes, then washed with 2 channel volumes of pre-warmed GMPCPP buffer to prevent nucleation from depolymerized dynamic microtubules, then incubated for 10 minutes and washed with 2 channel volumes of imaging buffer.

Drug-stabilized microtubules were bent by overflowing the channel volume during the incorporation step and alternating the direction of the flow by drying the channel with filter paper or by holding the microscope holder vertically, alternating either side of the channel for the duration of the incubation.

Recombinant His-DCX-eGFP-Strep (accession number NP_835365, Addgene #83918) was expressed in BL21(DE3) *E. coli* (New England Biolabs) at OD(600) = [0.4-0.6] and induced with 0.4 mM IPTG at 18 °C overnight. Cells were harvested by centrifugation, resuspended in Buffer A [50 mM NaPO_4_, 5 mM Imidazole, 300 mM NaCl, 5% Glycerol (v/v), pH 8.0] and lysed in an EmulsiFlex-C5 (Avestin). Cleared lysate was loaded onto a His60 Ni Superflow Resin column (Takara Bio) and eluted using Buffer A with a final concentration of 200 mM Imidazole and 10% glycerol (v/v). His-elution was subsequently diluted 1:5 into Buffer W [100 mM Tris-HCl, 150 mM NaCl, 1 mM EDTA, pH 8.0], loaded into Strep-Tactin Sepharose (IBA Lifesciences) and eluted with Buffer W supplemented with 2.5 mM D-Desthiobiotin and 10% Glycerol (v/v). Protein was loaded on a size exclusion chromatography column (Cytiva Superdex 200 Increase 10/300 GL). The peak fraction with the smallest mass (p) was recovered and concentrated. Protein aliquots were flash frozen in liquid nitrogen. Concentration was determined by absorbance at 280 nm with a DS-11 FX spectrophotometer (DeNovix) and purity was determined by SDS-PAGE (Fig. S5).

### Microtubule reconstitution imaging

Microtubules were imaged by TIRF microscopy on a Zeiss Axiovert Z1 microscope with a 100X /1.46 Plan-Apochromat objective (Zeiss, 420792-9800-000). Image acquisition was controlled using MetaMorph (Molecular Devices) and images were recorded on Andor Neo sCMOS or Photometrics Prime 95B sCMOS cameras with pixel sizes of 63 nm and 107 nm, respectively. Micrographs were taken at an exposure of 1 s to increase SNR and time-lapse micrographs were taken at an exposure of 100 ms. Afterwards, micrographs were processed in Fiji by Gaussian-filtered pseudo-flat-field correction, background-subtraction, and drift correction (Schindelin *et al*. 2012).

### Computational analysis

#### Curvature versus intensity plots

Quantification of fluorescence intensity and microtubule curvature were performed in Kappa, a Fiji plugin used to automatically fit the intensity of a microtubule to a B-spline curve (Mary and Brouhard 2019). In cells, images of single cells were taken with the Leica confocal microscope (see above). Microtubule segments were selected for analysis if they did not cross other microtubules and if they remained in the focal plane. *In vitro*, microtubule segments were selected if they measured more than 5.5 µm and their ends did not move between buffer exchanges. These segments were drawn by hand in Kappa and fitted to curves automatically using B-splines. These curves had an output of eGFP intensity (DCX), mCherry, Atto 550, or Atto 633 intensity (tubulin), and point curvature. In cells, intensity data was corrected by background subtraction. Corrected intensities were then normalized within each cell.

#### DCX phenotype classification

Images were taken with the OperaPhenix high content screening microscope to eliminate cell selection bias (see above). Regions of interest were drawn in Fiji around each visible cell to obtain a mean DCX-eGFP intensity. The DCX-eGFP phenotype was then classified as curves, puncta, or bundles.

#### Tip versus lattice intensity

Live cell images taken with the Lecia confocal microscope (see above) were selected where DCX-eGFP showed both a punctate and curvature localization within the same cell at 90 nM paclitaxel. Microtubule segments with a single DCX-eGFP phenotype (puncta or curve) were traced in Fiji to obtain a mean DCX-eGFP intensity and mean tubulin-mCherry intensity for the given segment. These intensities were corrected by background subtraction, and each segment was classified according to its phenotype.

#### Buckling assay

Quantification was performed in Kappa (Mary and Brouhard 2019). Microtubule segments were selected if they measured more than 5.5 µm and their ends did not move between buffer exchanges. The segments in the buckled conformation were drawn by hand in Kappa and fitted to curves automatically using B-splines. The fitted curves were re-fitted manually on to the straight conformation without moving the end points. Length measurements were output from Kappa and used to calculate the percent expansion relative to the length of a microtubule in the absence of drugs or DCX.

Paclitaxel titration expansion data were fit to a Hill equation

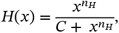

where n_H_ is the Hill coefficient, and

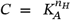

with K_A_ the half-saturation concentration and E_max_ the saturation value. The fit was made with the Python package “scipy.optimize.curve_fit” with “sigma” set as the standard deviation (Virtanen *et al*. 2020). The standard error of K_A_ was calculated with error propagation of n_H_ and C.

*Statistical analysis*. Statistical tests were Welch’s t-tests or Wilcoxon’s sign-ranked test, one or two-sided as indicated in figure legends. P values were considered nonsignificant when above 0.05

---

**Figure S1.**
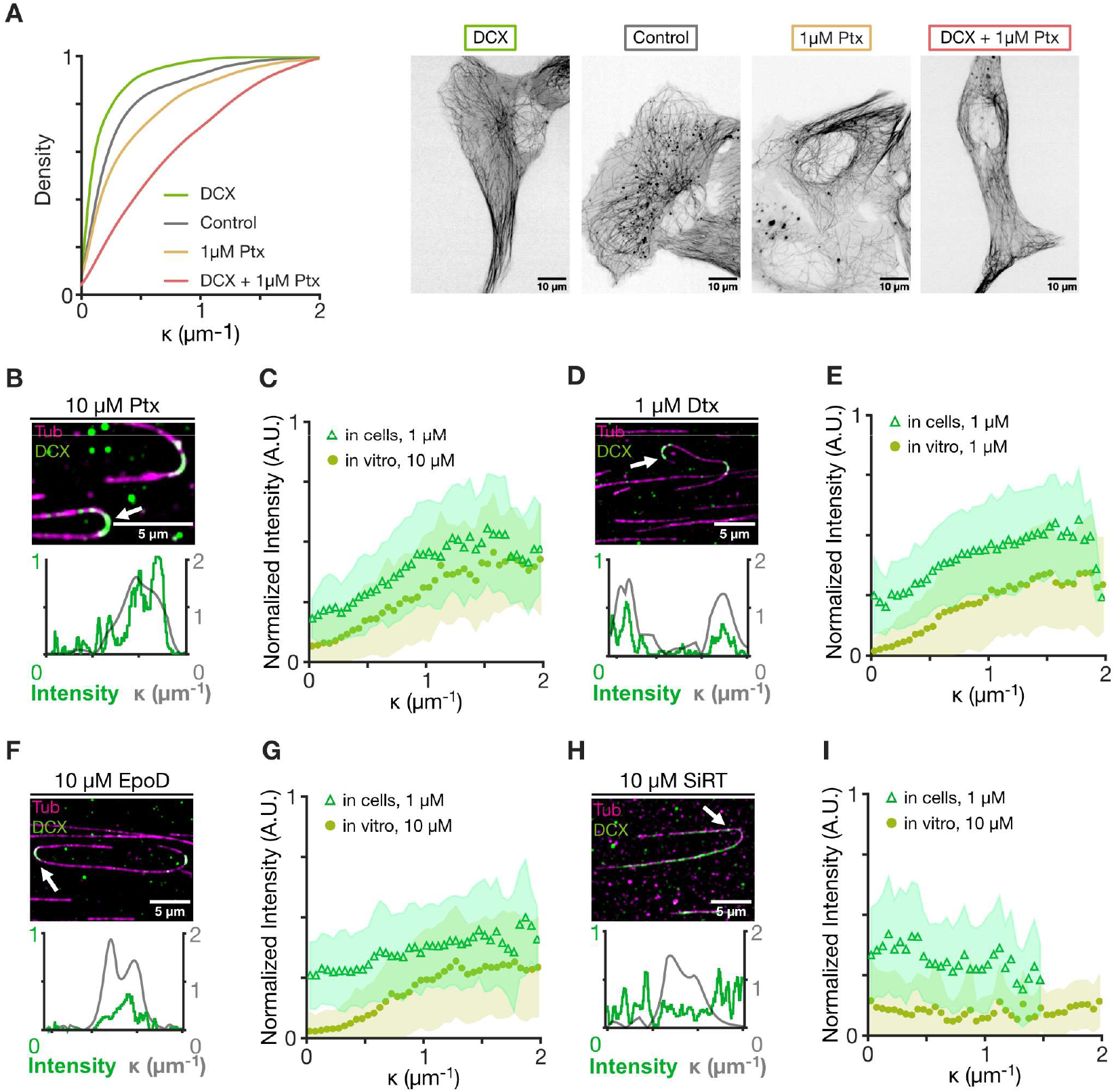
**(A)** *Left*: Cumulative distributions of microtubule curvatures. *Right*: Representative images of U2OS cells expressing Tubulin-mCherry corresponding to curvature distributions. **(B), (D), (F), (H)** *Top*: Micrographs of 5 nM DCX-GFP (green) binding to microtubules (magenta) stabilized with 10 µM paclitaxel (B), 1 µM docetaxel (D), 10 µM epothilone D (F), or 10 µM SiR-tubulin (H). White arrows indicate microtubules chosen for representative line scan. *Bottom*: Representative line scans of indicated microtubules. DCX intensity (green) and microtubule curvature κ (grey) are plotted over the length of the trace. **(C), (E), (G), (I)** Normalized fluorescence intensity of DCX-GFP in U2OS cells (light green triangles, data from Figs. 1G, 2B, 2E, 2H) and in reconstitution experiments (dark green circles) plotted as a function of absolute microtubule point curvature κ. Treatment concentrations are indicated on plots. n = 169 microtubules gathered from at least 15 micrographs and pooled across 3 repeats (C); 170 microtubules pooled from at least 15 micrographs across 3 repeats (E); 169 microtubules pooled from at least 15 micrographs across 3 repeats (G); 71 microtubules pooled from at least 15 micrographs across 3 repeats (I). Datapoints were binned and averaged in intervals of 0.05 μm^-1^. Shaded area represents standard deviation of the mean of each bin. See Table 1 for *p* values.

**Figure S2.**
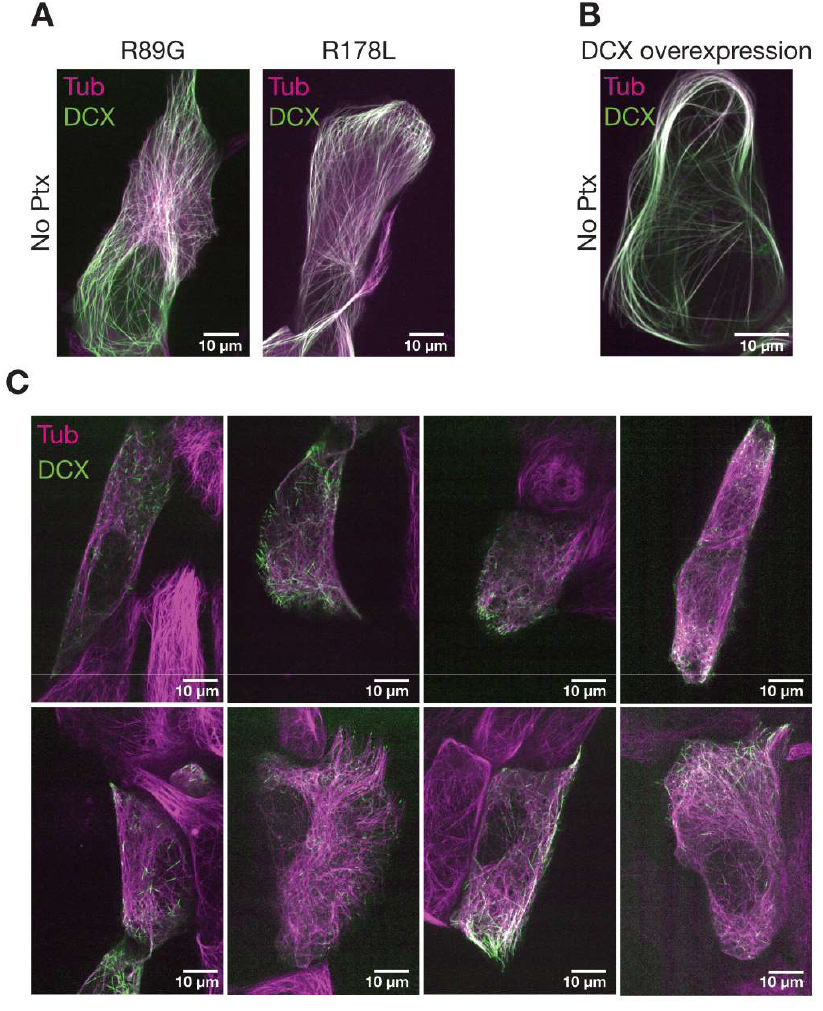
**(A)** Live cell images of U2OS cells stably expressing α-tubulin-mCherry (magenta), transfected with DCX-eGFP mutants (green): R89G (left panel) and R178L (right panel), without paclitaxel treatments. **(B)** Live cell image of a U2OS cell expressing α-tubulin-mCherry (magenta) and DCX-eGFP (green) and displaying a DCX overexpression phenotype. **(C)** Gallery of live cell images of U2OS cells expressing α-tubulin-mCherry (magenta) and DCX-eGFP (green) and treated with 90 nM paclitaxel.

**Figure S3.**
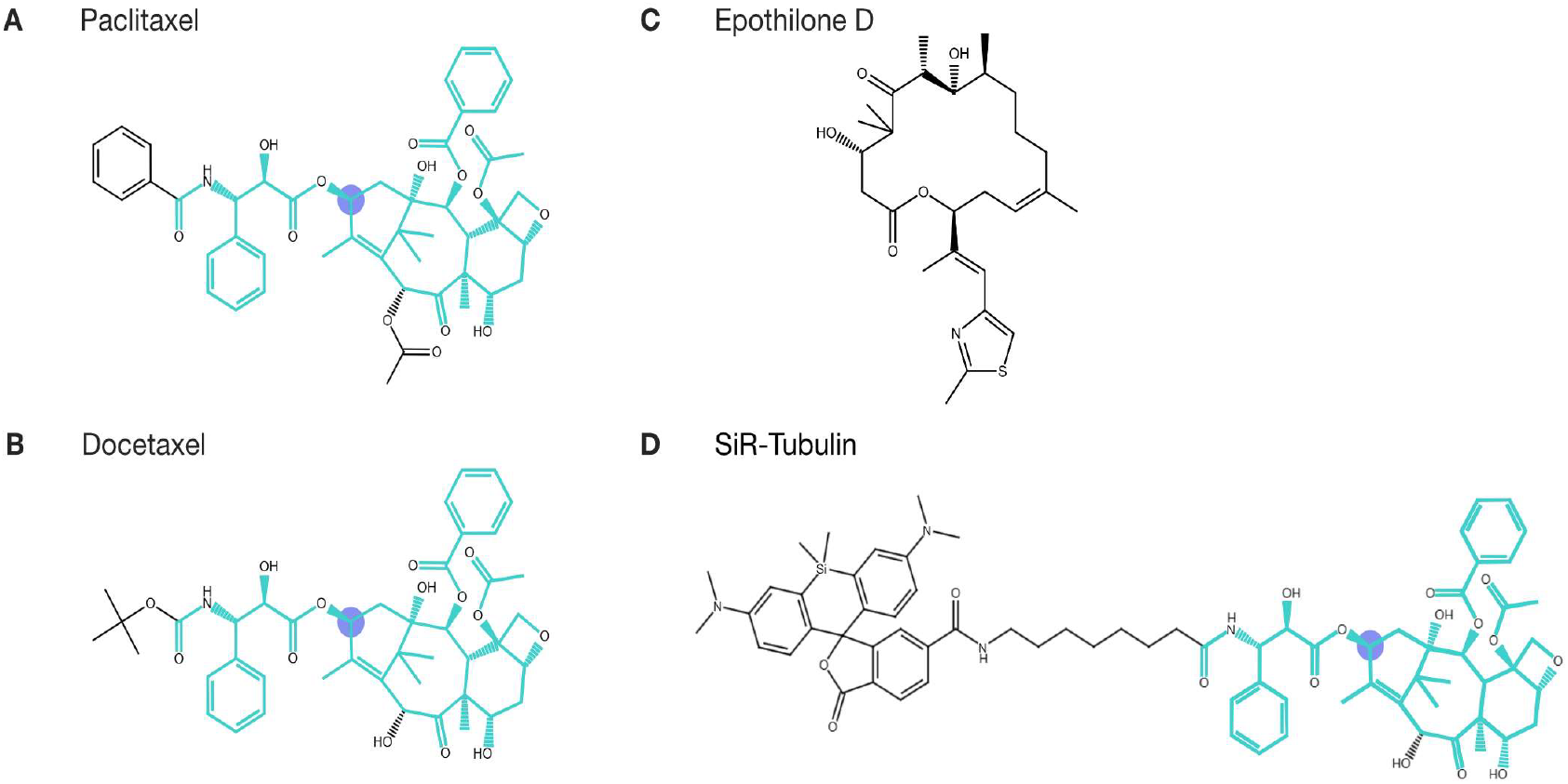
Chemical structures of **(A)** paclitaxel (PubChem ID 36314), **(B)** docetaxel (PubChem ID 148124), **(C)** epothilone D (PubChem ID 447865) and **(D)** SiR-tubulin (Lukinavicius 2014). The common core taxane structure common to paclitaxel, docetaxel, and SiR-tubulin is highlighted in cyan. The C13 side chain is highlighted in purple. Structures were generated in the RCSB PDB Chemical Sketch Tool.

**Figure S4.**
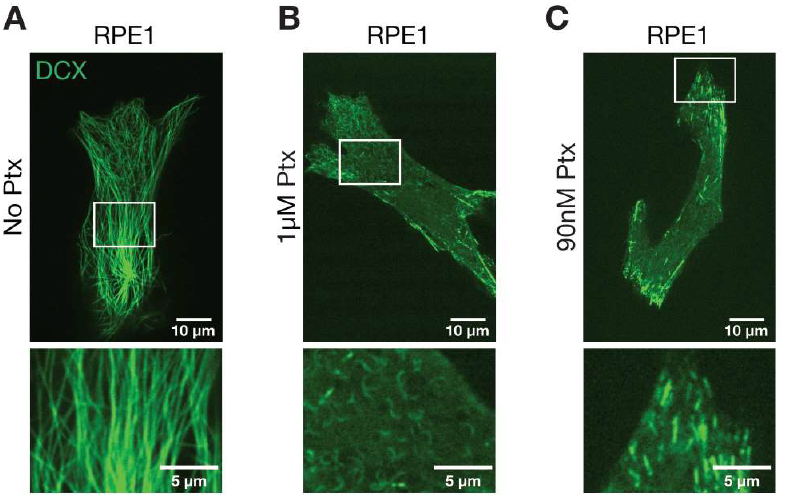
Live cell images of RPE-1 cells expressing DCX-eGFP (green). **(A)** Untreated. **(B)** After treatment with 1 µM paclitaxel. **(C)** After treatment with 90nM paclitaxel.

**Figure S5.**
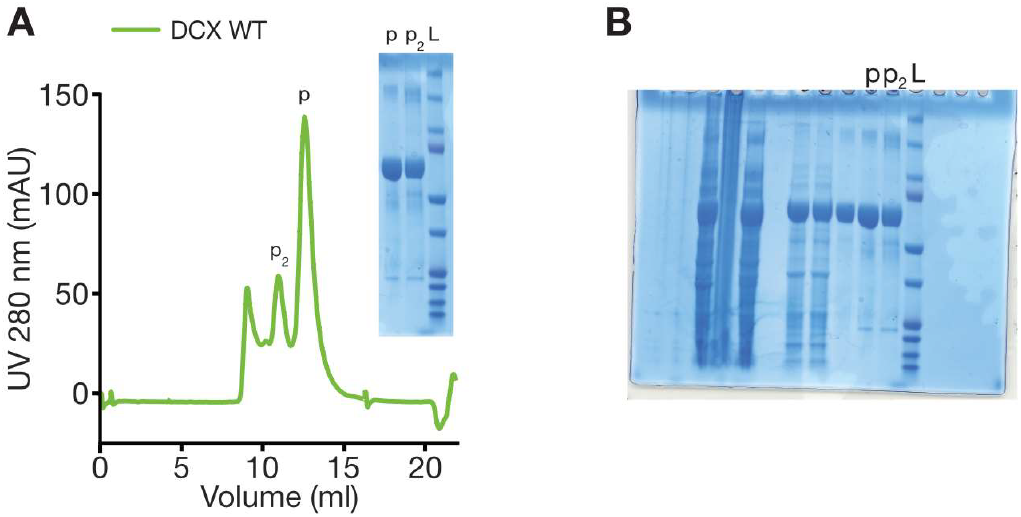
**(A)** Size exclusion chromatography trace of DCX-eGFP purification corresponding to SDS-PAGE gel of the peak fraction with the smallest mass (p), following the secondary peak (p_2_) and ladder (L). **(B)** Uncropped SDS-PAGE gel.

## Footnotes

1 We chose the term “buckling” because the microtubules experience axial compression forces when the length of the GDP-lattice increases between the two fixed GMPCPP caps.

